# Temporal integration and active sensing of light in planarians

**DOI:** 10.1101/2025.03.27.645397

**Authors:** Rimple R Dalmeida, Aalok Varma, Abhishek N Anil, Dasaradhi Palakodeti, Akash Gulyani

## Abstract

Animal light sensing and processing are intimately connected to the design of the sensory apparatus, with eye-design a subject of longstanding interest. Understanding animal response to stimuli can elucidate the nature of neural-processing encoded within the photosensory system, tuned to the specific eye-design. Planarians show exquisite photosensitivity and remarkably diverse light-induced behaviors. Further, they are bilaterally-symmetric and possess prototypic cup-shaped eyes, considered an important landmark in eye evolution. Here, using quantitative analysis of behaviour, we address planarian photo-response to distinct presentation of light gradients. Notably, we find that as the nature and presentation of gradients are altered, planarians switch their navigation strategies. When subjected to side-illumination, planarians display negative-phototaxis with a predominantly angular trajectory which presumably allows the animal to maintain the intensity difference between their two eyes for orientation during taxis. However, when subjected to top illumination where light intensity difference between the two eyes would be minimal, animals are still negatively phototactic but the dominant trajectories tend to be directly away from light. Apart from spatial light-intensity sensing, planarians also appear capable of temporal light sensing; confirmed by their response to a spatially-uniform light source with intensity modified over time. Crucially data show that planarians may be using spatial and temporal light sensing to differing degrees depending on the stimuli. Further, when subjected to light illumination from below, planarians fully switch orientation and go belly-up, indicating an active sampling of light distinct from photo-aversive behaviour. Overall, our results show planarian light-sensing to be multilayered, dependent on distinct, highly plastic sensory strategies and beyond basic photo-aversive behaviour.

## Introduction

All motile organisms need to detect and respond to gradients of sensory cues, in order to seek nutrition, a favourable environment or to avoid predators. Interestingly, a set of universal and overlapping strategies have evolved in nature to *detect* AND *respond* to the gradients. These strategies are in tune with the body size, body plan, overall physiology and more specifically the sensory and locomotive abilities of the organism in question. For instance, while some organisms may use a relatively simple strategy like ‘biased random walk’ to sample gradients, others might use more sophisticated methods to achieve gradient navigation(Hill & Häder, 1997). These may include strategies like spatial and/or temporal integration of sensory input to guide their response(Hinnemann et al., 2010; Sawin et al., 1994). Apart from conserved methods of gradient sampling to detect external stimuli, animals also adopt locomotion strategies that allow them to remain in the vicinity of a preferred stimulus or move away from a non-preferred stimulus; these include ortho and klinokinesis, where rate of turns and speed are modified depending on the strength of the signal sensed(Fraenkel G. S. Gunn D. L., 1940). Understanding these universal gradient-sensing and response strategies requires an in-depth investigation of sensory behaviour. In fact, a detailed analysis of gradient sampling and orientation behaviour can be richly informative about processes that may take place in the sensory nervous system and shed light on the mechanism underlying decision making(Gomez-Marin & Louis, 2012; Humberg et al., 2018; Kane et al., 2013). Such quantitative analyses enable insights into evolution of sampling strategies, behavioural plasticity, structure-function correlations in sensory apparatus and evolution of neural processing. Critically, the interplay between ‘sensing’ and ‘response’ mechanisms that is necessary for an organism to navigate its environment can be addressed.

Vision, having independently evolved multiple times, emphasises the importance of light sensing for the survival and propagation of species (Nilsson, 2013). Therefore sensing and processing of *light gradients* is of critical importance for many species. Interestingly, the strategies that organisms may deploy to sense and process light gradients are intimately connected to the sensory apparatus(Brodrick & Jékely, 2023; Jékely et al., 2008; Sawin, E. P. et al., 1994). Notably though, a wide range of light sensing structures exist in nature, with eye evolution having long fascinated biologists. Therefore, it would be interesting to examine if a framework for light gradient sensing strategies may emerge. Intriguingly, ‘single-cell organisms that sense light’ on one hand and ‘high resolution vision’ on another, have both been extensively investigated. However, eye-types that lie between these extremes, although highly abundant in nature, are relatively understudied. Understanding these eye types will help us connect how sensory processing and behaviour have evolved along with structure. One category of a highly ubiquitous eye type is the prototypical simple pigment cup eye where a pigmented cup encloses photoreceptor cells. In such eyes, the pigment presumably shades the photoreceptors from incident light from certain directions, thereby enabling the animal to sense direction. Such eyes often get classified under ‘low-resolution vision’ and may offer valuable insights into important landmarks in eye evolution(Brodrick & Jékely, 2023). Interestingly, the broad photosensory molecules in a cup-shaped enclosure is often retained in eye evolution(Nilsson, 2009).

Planarians, often studied for exceptional regenerative ability, possess prototypical pigment cup eyes as their primary visual organs(Morgan, 1901; Nilsson, 2009; Reddien, 2018; Reddien & Sánchez Alvarado, 2004; Sánchez Alvarado, 2006). In fact, planarian light sensing captured the interest of early biologists, likely due to prototypical cup-shaped eyes and robust light sensitivity(Darwin, Charles, 1859; Mast, 1911; Morgan, 1901). More recent work has revealed planarians to be capable of remarkably complex and multifaceted light sensing behaviour and associated sensory processing, with light sensitivity even extending beyond the eye(Akiyama et al., 2018; Birkholz & Beane, 2017; Paskin et al., 2014; Shettigar et al., 2017, 2021). The planarian visual system appears capable of high-fidelity sensory processing, despite having a so-called ‘simple’ sensory apparatus and visual network, especially in terms of the number of cell types that form the eye and the lack of focusing structures. For instance, planarians, by virtue of having only one opsin in the eyes(Azuma et al., 1999), were thought to be ‘colour-blind’ or incapable of distinguishing light wavelengths. However, it was discovered that just a small change in wavelength of light (as little as 25 nm) can completely switch their behaviour in binary-wavelength choice assays(Shettigar et al., 2017). The remarkable light sensing abilities of planarians mentioned above as well as regeneration abilities make planarians an attractive model to interrogate light sensing(Atabay et al., 2018; Deochand et al., 2016; Inoue et al., 2004; Ivankovic et al., 2019; Owlarn & Bartscherer, 2016). The fact that planarians have a relatively simple eye type, a body plan with bilateral symmetry, presence of well-demarcated cephalic ganglia along with a delocalised nerve nets reminiscent of the cnidarian nervous system; adds to the attractiveness of addressing planarian light sensing. In fact, more broadly, light sensing in platyhelminths is relatively understudied, adding to the need to address this gap.

The sophisticated sensory processing demonstrated by planarians offer unique opportunities to study specific sensory strategies deployed by animal with a visual system having bilaterally organized simple cup shaped eyes(Inoue et al., 2015). Planarians are sensitive to small changes in light stimuli. The remarkable wavelength induced photoswitching in binary choice assays(Shettigar et al., 2017) appears to be due to sensing small changes in ‘effective’ intensity absorbed at the eyes. According to conventional understanding of wavelength sensing, wavelength discrimination emerges through distinct photoreceptors for distinct wavelengths. Organisms with just a single broad-range photoreceptor would *apriori* be insensitive to light wavelengths. However, planarians, despite having a single broad photoreceptor, appear to be acutely sensitive to the amount of light absorbed (‘effective absorbance’). When presented with two light inputs of wavelengths just ∼25nm apart (and thereby small differences in amount of light absorbed), these organisms show consistent choices in binary choice-assays(Shettigar et al., 2017). With this background, we surmised that planarians, apart from being highly aversive to light, are likely to be very sensitive to light gradients. Further, we hypothesised that planarian light-gradient sensing may be highly plastic and tuneable to the type of gradient. To address this, we have examined specific sensory strategies planarians deploy to navigate light gradients; if planarian sensory strategy is sensitive to the type of gradient presented and how?.

We have also addressed interplay of sensing with locomotion; since movement forms an intricate part of the response. Does response (movement) connect back to the sensory strategy and how? Planarian motility generally involves gliding on surfaces with a ‘wigwag’ movement of the head(Akiyama et al., 2018; Pearl, 1903; Taliaferro, 1920; Walter, 1907) where the sensory strategy they appear to employ is to move their heads and assimilate light stimuli. However, the range and breadth of sensory strategies planarians employ is unclear. Importantly, whether (and if yes, how) these organisms modulate and tune the sensory strategies to changing stimuli is unknown. Such information will not only help answer unaddressed questions specific to the planarian nervous system but also address the broader question of the capabilities of a supposedly rudimentary ‘prototypic’ visual system(Nilsson, 2013).

Here, we have performed a systematic analysis of planarian response when subjected to controlled visual stimuli. Towards this end, we designed light assays and automated tracking methods where multiple parameters of planarian sensory behaviour and locomotion can be recorded. Crucially, the light assays and stimuli have been designed in a manner so as to ask how sensory and response strategies may depend on the structure of a simple bilaterally symmetric visual system. For instance, the assays were designed to modulate the difference in intensities sensed at the two eyes. Specifically, we compared responses to light gradients when the difference in the intensities sensed at the two eyes were expected to be large versus a scenario where the difference in intensities may be significantly smaller (See Figure 1 for design). The source of light was positioned in specific positions to achieve this. We hypothesized that this design would help us look deeper into planarian visual processing capabilities, and the type of strategies deployed. Notably, we found that planarians may indeed be deploying multiple strategies for sensing light gradients. In a scenario where there may be substantial differences in the light intensities sensed at the two eyes, planarians appear to be relying on ‘spatial’ sensing of gradients. Moreover, we discovered that the trajectory of locomotion response appears to *further* augment the spatial sensing strategy by ensuring that the spatial differences between light intensity at the two eyes is maintained when animals move away from light. Remarkably, however, the type of gradient significantly impacts the sensing strategy of the animal and nature of response. If spatial differences in light intensity are not large, locomotion trajectories and sampling behaviour are completely altered with planarians likely relying on *temporal* processing of light stimuli to navigate gradients. Temporal processing was also confirmed by designing assays with uniformly lit arenas, modulated over time. Therefore, we show planarians capable of both spatial comparisons as well as temporal integration of stimuli, with the contribution of each strategy depending on the nature of stimuli. Intriguingly, we also found that planarians while clearly being acutely light aversive, actually sample and move to locate the source under certain unrelenting light exposure, again suggesting a surprisingly active sampling of light stimuli rather than passive/monotonic movement away from ‘aversive’ stimulus. This study reveals remarkably active and highly plastic light sensing and adaptable navigating strategies employed by planarians, and sheds light on how even a bilaterian with an relatively uncomplicated visual network may navigate changing light conditions and complex gradients with a stunning array of sensory responses.

**Figure 1:**
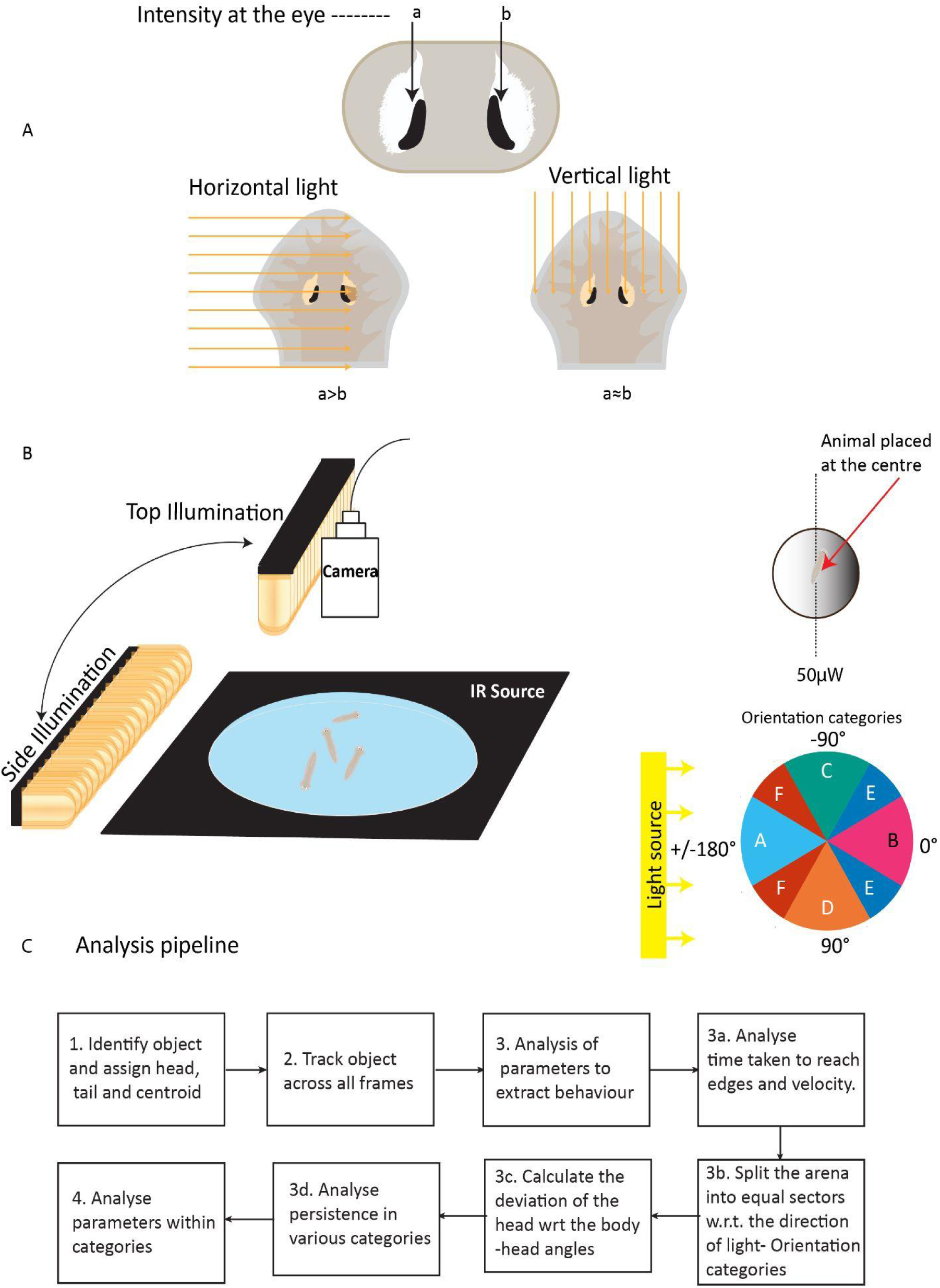
Rationale behind the assay design and the analysis of phototaxis A) Illustration of the incident light under the two conditions depicting the difference in light intensities between the two eyes. Intensities at two different eyes are considered to be a and b. In horizontal light condition a can be greater than b; in vertical light condition a is similar in value to b B) Illustration of the LED panel positioning with respect to the arena and the camera, IR source is under the circular arena to illuminate it for the camera(left). Location of the power measurement at the arena(right). C) Orientation categories are defined with 4 categories of 60 degrees each, A,B,C, and D as shown. The categories E and F are 30 degrees each and are symmetrically placed facing away and towards the light centred at +/-45 degrees and +/-135 degrees. D) Flow chart depicting the analysis pipeline and the parameters studied. The key stages of the pipeline involves identifying the object/animal and assigning head and tail coordinates along with the centroid. This is followed by tracking the position of these three points on the animal across all frames of the recording. Using the data from the tracking, speed, orientations, head angles and persistence are analysed as described in Materials and Methods section.

## Results

### Designing distinct presentation of light gradients and assay designs

We wanted to examine light gradient sensing in planarians and ask if worms are sensitive to the type of gradient presented. Based on past work(Shettigar et al., 2017), we surmised that worms may be able to distinguish light intensity differences between their two eyes for complex sensing and navigation. It appears that the structure and organisation of the planarian eyes with the presence of an optic chiasm may facilitate differential intensity sensing between two eyes(Fumie Sakai et al., 2000; Okamoto et al., 2005). Planarian eyes are made of laterally facing pigment cups housing the photoreceptors with a slight anterior tilt which can provide directional information to the animal(Akiyama et al., 2018; Carpenter et al., 1974). If light is incident horizontally, the organisation of the eye will illuminate the proximal eye to a greater extent than the distal eye at most instances except when the animal is positioned parallel to the direction of the incident light. Therefore, we estimated that side illumination (SI) or ‘horizontal’ light inputs would tend to maximize the difference in intensities between the two eyes (Figure 1A). On the contrary, top illumination (TI) or vertical light would provide less intensity differences between two eyes. Therefore we hypothesised that analysis of behaviour under these contrasting conditions could provide insights in light sensing mechanisms.

The source used was made of a single white LED panel to achieve even illumination. The Illuminated area was wider than the arena to prevent any unilluminated regions. The two assays namely side illumination (SI) and top illumination (TI) assays were designed as shown in Figure 1B. The intensity at the point where the animal was dropped was maintained at 50 µW/cm^2^ in both assays. An IR light source enabled videography. Videos capturing worm behaviour and movement recorded were analysed using programs written in MATLAB as described in the methods section under ‘analysis pipeline’. The analysis was stopped when the worm reached close to the edge of the arena as they tend to show a preference for edges(Akiyama et al., 2015). Key steps for video analysis are outlined in the flowchart in Figure 1D.

In order to understand the sensory behaviour and the potential underlying strategies, we examined the orientation of the animals (orientation angles) with respect to the source of light. Orientation angles were measured by calculating the angle between the direction diametrically opposite the light source and the direction the animal is headed towards (orientation) at any given point. For instance, if an animal heads directly away from the light source, orientation angle would be zero (see Figure 1C). Direction of movement or orientation of the animal is represented by the vector passing through the tail and the centroid(Supplementary Figure 1, also see methods section). Orientation vector used here is distinct from the vector from the centroid to head. ‘Centroid to head vector’ may be more variable and subject to sampling noise due to the natural wigwag movement of planarian heads(Akiyama et al., 2018). In order to resolve the orientation angles meaningfully, the orientation angles in the circular arena were split into equally weighted six categories of sixty degrees each (Figure 1). Categories A, B, C and D consisted of continuous angular cones spanning sixty degrees each; while two symmetrical angular cones spanning thirty degrees each together composed each of categories E and F. Category A had orientations directed towards light while category B had orientations most away from light, each spanning cones of sixty degrees. C and D categories were centred around orientations perpendicular to the light source. E category split into two symmetrical orientation cones of thirty degrees, each cone of thirty degrees centred at +/-45 degrees (pointing away from the light source). F category split into two symmetric thirty-degree cones, each cone centred at +/-135 degrees towards the light source. The orientation categories allowed binning of orientation of worm movement in response to light gradients, enabling further analysis of gradient sensing behaviour and underlying navigation strategies.

Apart from overall worm orientation, other parameters of the animals response to light gradients were quantitatively examined. One of them was ‘head angle’; defined as the angle subtended by the head with respect to the overall orientation of the animal. Head angle helps describe the natural wigwag motion of the head, a feature commonly associated with planarian motility and has been examined before (Akiyama et al., 2018. Two other parameters examined here are the ‘speed’ and ‘persistence’ of movement. Persistence of movement, studied to examine presence of persistent runs, is defined as a specific distance covered by an animal in the forward posture within +/- 10 degrees before a sway, which in itself is defined as a head angle change larger than +/- 10 degrees. All of these parameters provide a comprehensive quantitative investigation of planarian response to light gradients and navigation strategies.

### Planarians switch trajectory as stimuli presentation is altered

Using the design and methods outlined in the previous section, animals were subjected to light gradients generated by either side (SI) or top (TI) light illumination and the behaviour analysed. Notably, we found that the precise presentation of gradient has a clear impact on animal behaviour and movement trajectory, with animals completely switching their trajectory patterns as the gradient presentation is altered. When animals were subjected to top illumination, the trajectory of movement tended to be directly away from light. This can be seen from a 2-D plot of worm coordinates (Figure 2A), where a clear clustering of animal trajectories away from light can be observed. This movement of worms away from light is also illustrated by an analysis of orientation angles (Figure 2C). A histogram of the orientation angles shows a clear unimodal distribution away from light. On the contrary, the behaviour of animals subjected to light gradients generated through side illumination (SI) is bimodal and hence characteristically different. 2-D plot of coordinates depicting worm trajectories shows a striking ‘split’ in the distribution. Interestingly, under SI, worms appeared *not to populate* trajectories directly away from light, unlike what was seen with top illumination. Worms appeared to consistently move along distinct angular trajectories, while showing negative phototaxis. The distribution of orientation angles reinforce the contrast between SI and TI. Orientation angles directly away from light tend to be under-sampled in SI assays, distinct from the behaviour in TI. A histogram of orientation angles under SI shows a clear bimodal distribution, again suggesting clear, discernible angular movement for negative phototaxis. This is consistent with a report of a split in distribution of orientation angles during phototaxis in *Dugesia japonica* (Akiyama et al., 2018). Notably, however, we find that this observed angular movement seen in SI, illustrated by both worm trajectory and orientation, is extremely sensitive to gradient presentation, and is not observed in TI.

**Figure 2:**
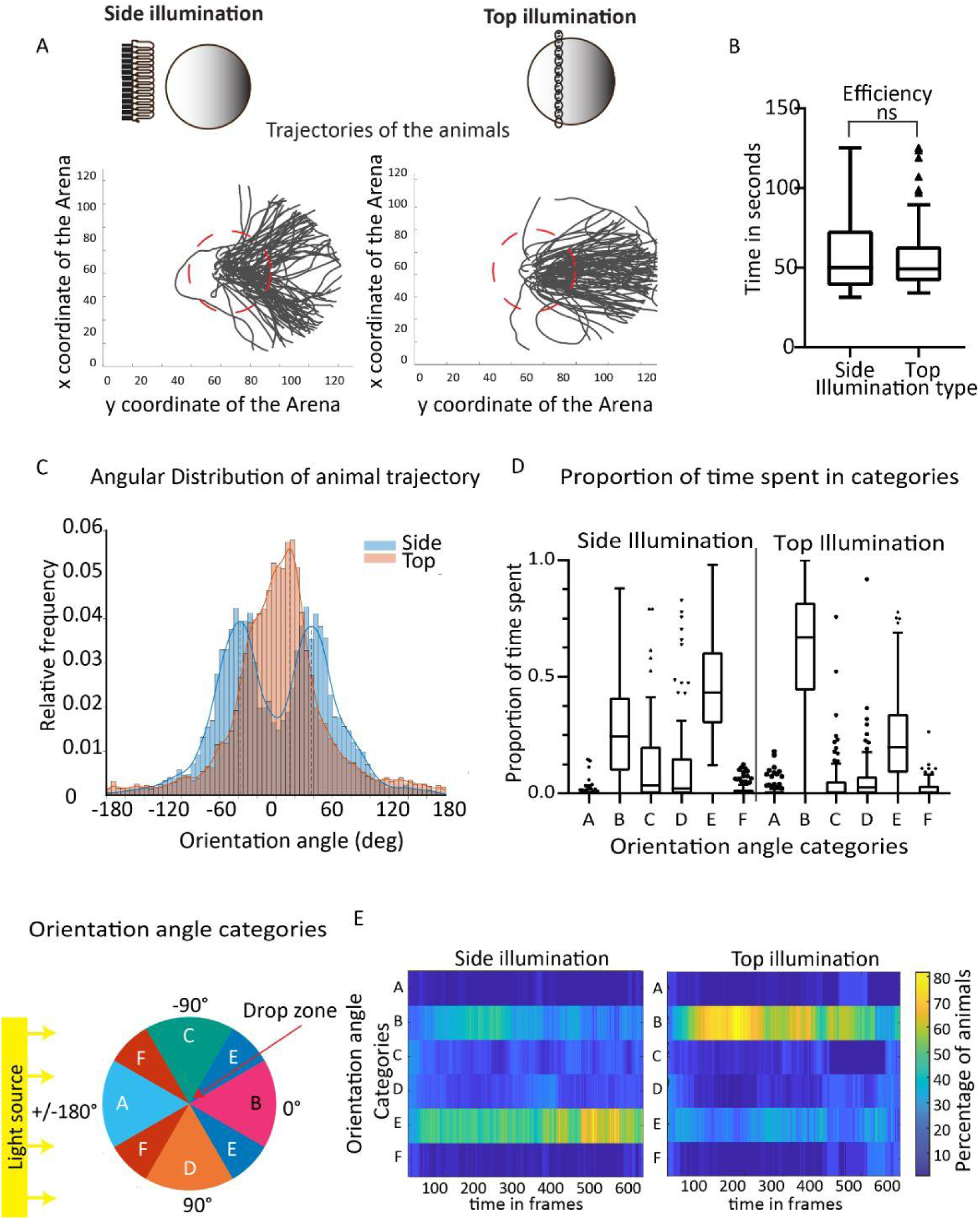
Versatile phototactic response to the two assays (A)Trajectories followed by the animals on the arena over time in the SI and TI assay after being placed in the drop zone shown in red dotted circles and observed for 2 minutes. (B) Time taken to reach the end of the assay under SI and TI conditions. (C) Histogram of orientation angles in side(blue) and top(red) illumination assays. (D)Proportion of time spent in the various orientation angle categories, defined in the legend at the bottom left of the panel, under the two conditions (N= 89). (E) Heatmap of the proportion of animals in the orientation angle categories represented as a percentage over time. Each horizontal band is a category and the x axis represents the frames during the course of the assay where a frame is captured every 150ms. Colour coding shows the percentage of worms in each category at a particular time, as shown in the colour key. ns p>0.05

The proportion of time spent by a worm in various orientation categories (Figure 1D3b and Figure 2D) as they navigate gradients is also instructive. Worms preferentially populate certain orientation categories, under both side and top illumination. However, the orientation categories populated are clearly different in the two assays. Under TI, worms appear to rapidly populate orientation category B, composed of orientation angles directly away from light. However, under SI, worms appear to orient preferentially at angles about +-45 degrees relative to direction directly opposite the light source (category E). Some animals are also seen along orientation angles directly away from the light source. Orientation appears to occur faster for the TI compared to SI, which appears to be more gradual (Figure 2E).

Interestingly, while the sensory responses and the trajectories used by the animals when subjected to top and side gradient appear very characteristic, the time taken by worms to under the two assays is comparable (Figure 2B). While many aspects of planarian movement under light stimulation have been studied before(Akiyama et al., 2018; Parker & Burnett, 1900), our data provides clear evidence that planarians appear to deploy multiple strategies to orient and navigate light gradients. Further, these underlying sensory strategies appear to be plastic and extremely sensitive to changes in the angle of gradient presentation.

### Multifaceted strategies include head-sways for gradient sampling

Our results so far indicate planarians switch trajectories as gradient presentation is altered, and appear to use their trajectories to augment sensing (see Figure 2). We further analysed the phototactic behaviour to identify other aspects of the behaviour that shed light on sensory strategies. Using the categories previously defined and splitting the dataset into animals moving away and towards light allows us to isolate the speed at which they move in these categories (Figure 3A). We found that animals move slower when moving towards light compared to when they move away. Such response is suggestive of orthokinesis(Gomez-Marin & Louis, 2012), wherein the animal modifies its speed based on the gradient of stimulus. Notably, this is more pronounced in SI assay compared to TI assay (Figure 3A).

**Figure 3:**
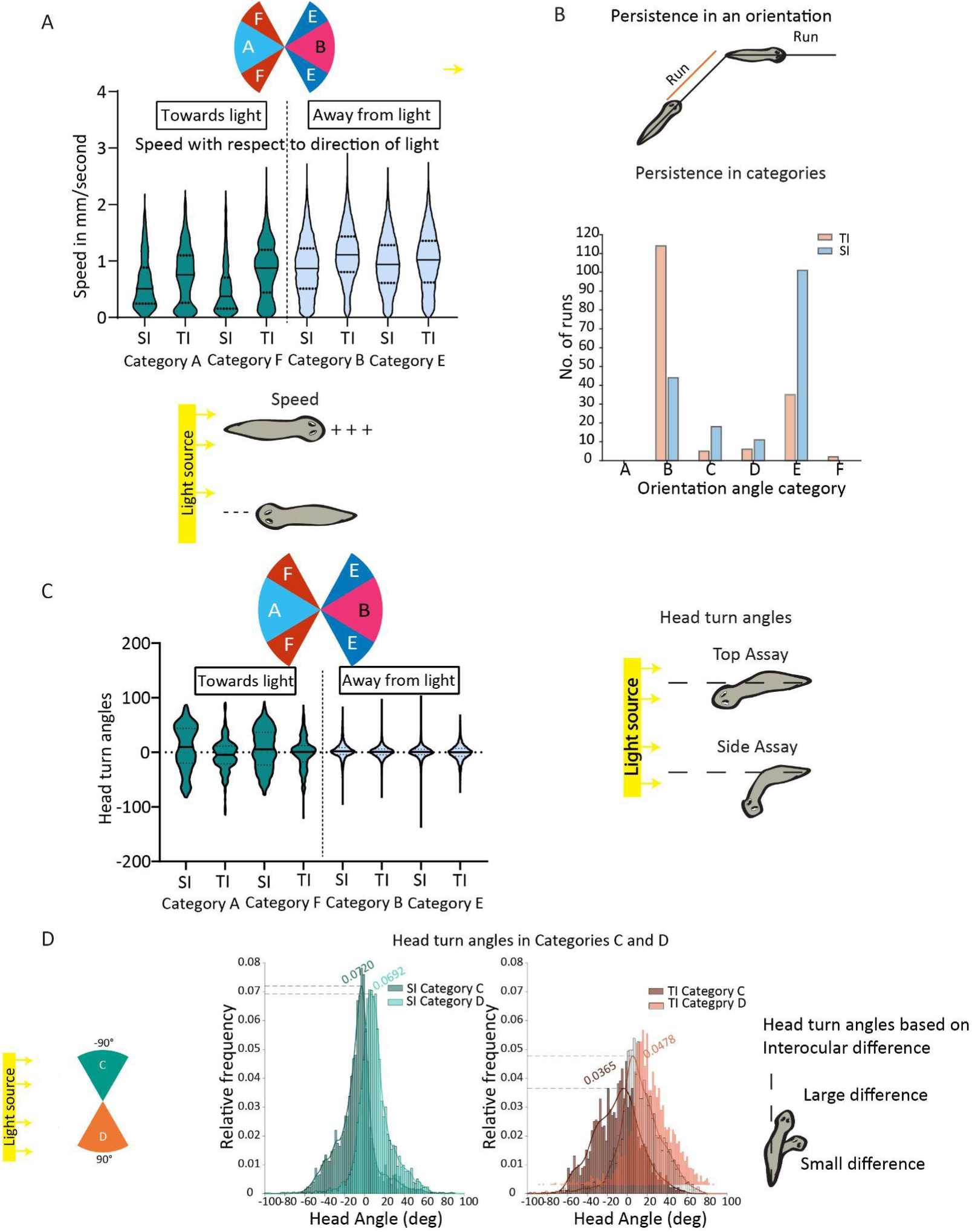
Temporal and spatial processing utilised for fine gradient sensing (A) Violin plots of speed grouped in conditions going towards and away from light. (B) Number of runs in various orientation categories with SI(red) and TI(grey) assays. (C) Violin plots of head angles grouped in conditions going towards and away from light.Levene’s Test results: F-value, p-value 123.01, p<0.0001 [SI vs TI, A];F-value, p-value 110.46, p<0.0001 [SI vs TI, F] (D)Head angle histogram in C and D categories for SI(left) and TI(right) assays. A chi-squared test is used to compare the proportions of worms in each category as a function of the assay. The null hypothesis assumes that the proportion doesn’t depend on the assay being used. The chi-squared statistic obtained is 73.5257, with 4 degrees of freedom (# of cols in tbl - 1)x(# of rows in tbl - 1). This produces a p-value of 4.0847e-15, which means that the null hypothesis can be rejected in favour of the alternate hypothesis.

Apart from speed, light also alters the persistence of movement. Persistence is analysed by examining the number of ‘runs’ made by the animal. A ‘run’ is defined as movement for a certain fixed distance (1.5 mm) with head turn angles not crossing a threshold of ten degrees (Figure 3B). Data show that for both TI and SI animals tend to have more ‘runs’ away from light than in other directions. This reduction in ‘runs’ towards the light source suggests that planarians may sense an increase in intensity over time, rather than rely exclusively on spatial sensing and initiate turns as light intensity increases. The contribution from a potential temporal sensing would increase as the animals would move either completely towards or away from light.

We then examined if the animals use additional ways to sample distinct light gradients at any location. For this we examined head angles subtended by the animals when subjected to top and side gradients. Figure 3C shows that irrespective of the type of gradient presented, the measured ‘head turn angles’ are more broadly distributed when moving towards the light compared to when animals move away from light. This indicates increased sampling as the light intensity increases, and further suggests that animals are able to sense how light intensity changes over time. Therefore, apart from spatial sampling (intensity difference between two eye spots; see previous sections), animals may also deploy temporal integration of light intensity sensed at the eyes. Such temporal integration would allow an animal to distinguish between whether the light is increasing or decreasing as a result of movement.

While the spread in head turn angles consistently increases as animals move towards light, there are interesting differences between top and side illumination scenarios. The spread in head turn angles during movement towards light is much greater under SI versus TI (Figure 3C; comparing categories A, F *versus* B, E). It would appear that animals, when moving towards light under SI, sample more (through head turns) in order to maximize difference in light intensities sensed by the two eyes. Increased angular sampling is manifested through larger head turn angles seen under SI (Figure 3C).

Subtle differences in sampling behaviour under SI vis-à-vis TI can also be seen in the analysis of head turn angle in categories D and C i.e. when animals are moving at right angles to the light gradient. In these orientations, one would expect the greatest difference between intensities sensed at the two eyes under SI whereas the contrast would not much less in TI. This is reflected in the spread of the head turn angles. Interestingly, under TI animals tend to sample light through large head turns when they move perpendicular to the light (Figure 3D); whereas under SI smaller head turn angles tend to be observed (categories D, C). It would appear animals rely on larger head turns when moving perpendicular to the light in order to overcome the challenge of lesser contrast between the eyes under TI. These results again illustrate how animals can fine-tune or even switch their sensory and locomotion strategies as they navigate distinct stimuli.

### Response to temporally changing uniform light indicates temporal processing

The results so far show remarkable plasticity in planarian sensory behaviour, including the possibility of temporal sensing coexisting with spatial sensing strategies. We examined the potential temporal sensing capabilities further by studying the response of the animals to a uniform light environment that changes over time (no spatial variations, exclusively temporal changes). This was accomplished by uniformly illuminating an arena from above the animals and automatically changing light intensity over time (Figure 4A).

**Figure 4:**
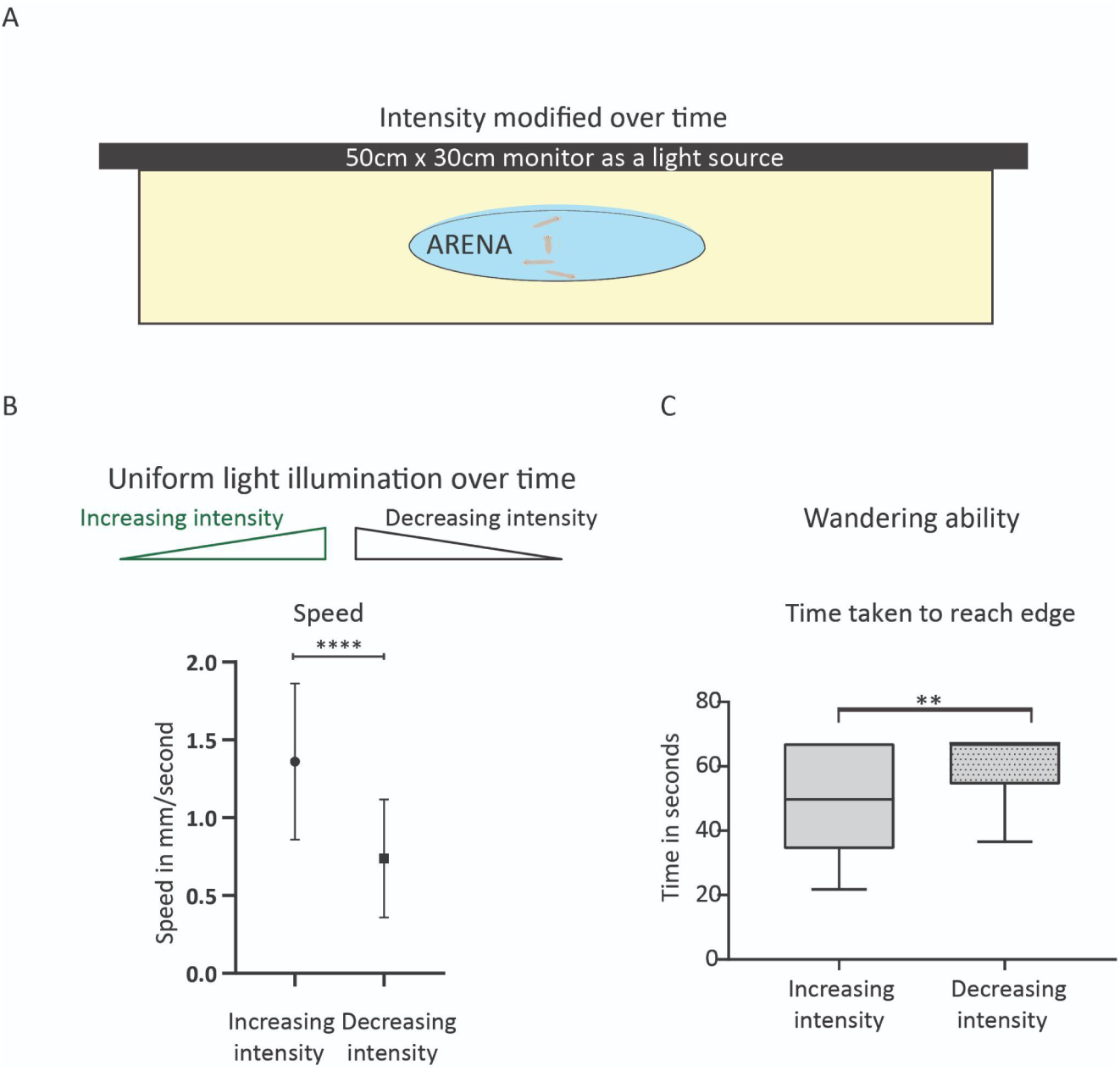
Uniform light assays for understanding temporal processing. (A)Assay set up with the arena placed under a monitor with intensity modified uniformly, over time. (B) Speed of animals under increasing and decreasing intensity. (C)Time taken by the animals to reach the edge plotted as box plots. N=62; **p<0.01, ****p<0.0001 Welch’s t test

We find that in the uniformly lit arena, the speed of movement of the animal under increasing light intensity is significantly higher than when the light intensity decreases over time (Figure 4B). These data suggest animals are able to sense changing light conditions over time and modulate their behaviour, even when there are no spatial gradients. It would appear that on decreasing light, photo-aversive planarians sense more ‘favourable’ light conditions and reduce speed. Such a sensory strategy would presumably allow the animals to remain in the vicinity of lesser light. On the contrary, animals speed up when light intensities increase over time, indicating an enhanced escape response.

These findings are further strengthened by examining the time taken by the worms to reach the edge of the arena as light intensities change over time(Figure 4C). Notably the time taken to reach the edge tends to be much shorter when the light intensities increase over time as compared to time taken under decreasing light intensities. These data reinforce the idea that animals are able to clearly sense increases in light over time, and in response to that show an escape like behaviour, rapidly reaching the edge of the arena. Together the data suggest planarians do indeed have the ability to integrate the amount of light sensed over time (temporal integration of light intensities sensed by the eyes) and modulate their behaviour accordingly.

### Light dependent ‘belly-up’ flipping highlights sensory strategy diversity

Planarians rapidly tune their sensory and response strategies, and appear to acutely sample light in diverse ways. Remarkably, here we also found that planarians rapidly flip their dorso-ventral orientation in a light-dependent manner. When an arena is uniformly illuminated from below(Figure 5A), we observed that planarians tend to flip to a belly-up orientation, clinging on to the surface of the water (Figure 5B). Notably, this belly-up behaviour was not seen to any significant degree when worms are illuminated from the top. This behaviour was also not observed in the absence of light for the duration of the assay.

**Figure 5:**
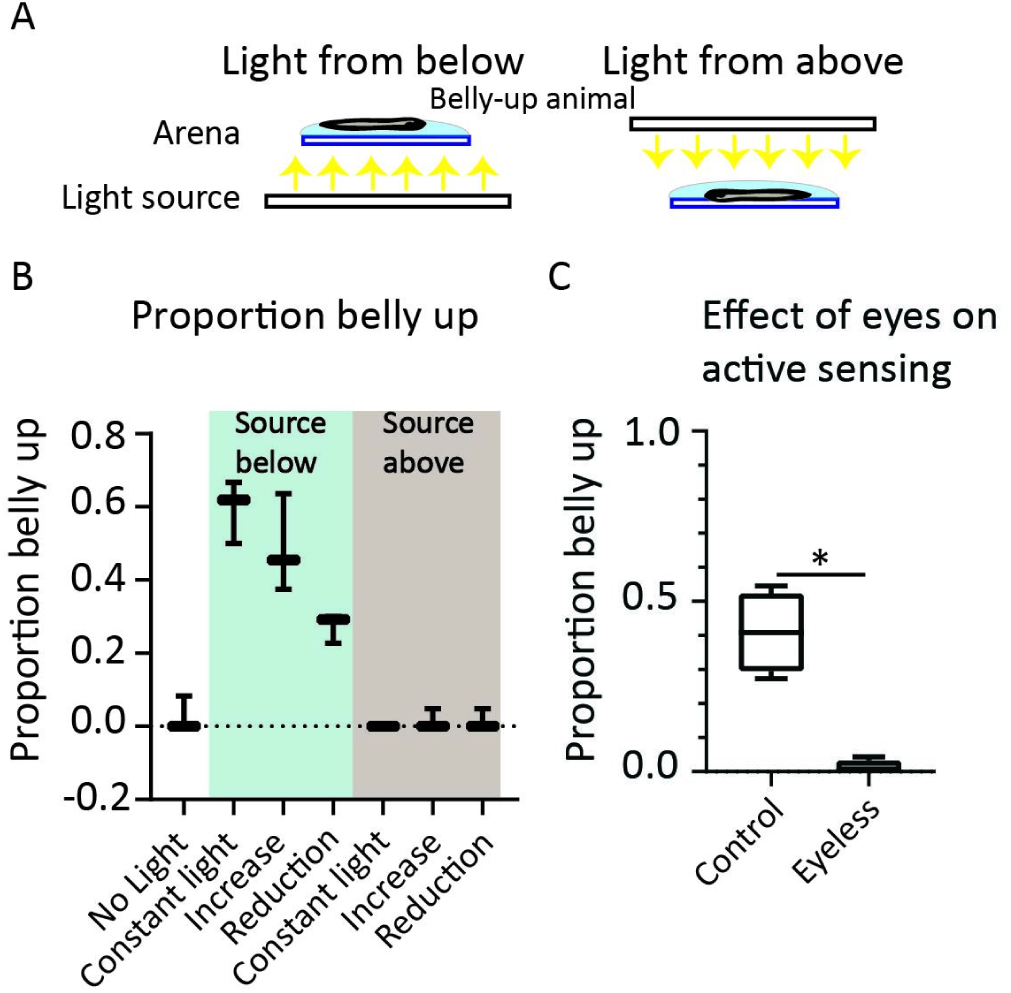
Active sensing under uniform light conditions A) Panel shows the light set up with the arena illuminated from below and above. B) Proportion of animals belly up under the conditions specified on the x axis. The pale blue is where the light is from below and the brown is where the light is above. The first box plot is under ‘no light’ condition. C) Boxplot of proportion of animals belly up under control and eyeless condition. (B: N= 68; C: N= 88) ; *p<0.05, **p<0.01, ***p<0.001

Further, this orientation flipping is dependent on changes in light intensities. More orientation flipping is seen when light intensity increases over time, as compared to decreasing light intensities (Figure 5B). We then examined if the orientation behaviour is dependent on the visual system. For this we generated eye-less animals through a RNA-i-mediated knockdown of the transcription factor ‘ovo’. Animals were amputated and during head-regeneration, animals were injected with ds-RNA targeting ovo transcript in order to generate eyeless animals(Cross et al., 2015; Lapan & Reddien, 2012). Previously eyeless animals following ovo knockdown were shown to fail to carry out phototaxis(Cross et al., 2015). Notably, belly-up orientation flipping is not observed in eyeless animals(Figure 5C), clearly demonstrating that this behaviour is dependent on visual light sensing.

These findings suggest that planarians, while being photoaversive, do not have just an indiscriminate and unrefined escape response. We interpret these findings to suggest that photoaversive planarians, while moving to minimize overall light exposure, also likely ‘actively sense and sample’ the presence and location of the light source/stimuli. Such a response was also seen in *Drosophila* larva (De Andres-Bragado et al., 2018; Gomez-Marin & Louis, 2012). Planarian eyes are located towards their dorsal side. When light is incident from below, flipping ‘upside-down’ would presumably allow the worms to locate the light source.

## Discussion and summary

This work demonstrates planarian light gradient sensing to be remarkably rich and multifaceted, indicating highly plastic and adaptable sensory strategies. By providing planarians with diverse light inputs and systematically recording and analysing planarian responses, many new behaviours and response patterns have come to light. A notable finding is that presentation of gradient i.e. the direction of incident light, can completely switch sensory behaviour and locomotion trajectories. In the scenario of ‘side illumination’, planarians show angular trajectories, while ‘top illumination’ leads to worms typically moving directly away from the direction of light. This finding points to planarians using subtly distinct sensory trajectories when navigating light gradients. An angular trajectory while moving away from light would likely mean light intensity sensed at each planarian eye would be different. Therefore, animals retaining intensity differences between the two eyes over the course of their movement away from light, may help worms navigate efficiently using a bilateral contrast strategy. This would be an example of spatial sensing, where intensity sensed at distinct points in space would need to be different. In this case the trajectory of movement during side illumination, augments and helps with spatial sensing. Angular trajectories during negative phototaxis have been reported for a distinct planarian species(Akiyama et al., 2018).

In contrast, if the gradient is generated through a ‘top illumination’ (lower intensity differential between the two eyes), majority of the planarians tend to move directly away from light. This may be due to the fact that spatial (bilateral) contrast is limited and relying primarily on spatial sensing may not be efficient. It appears in such a scenario the animals switch to a different strategy for navigating light gradient i.e. ‘temporal sensing’. Temporal sensing is understood to involve animals sensing signal (light) intensity over time and using this change in signal over time to navigate gradients. This is also referred to as temporal integration of signal. In this case, integration of light intensities would allow planarians to sense if the light intensity is increasing or decreasing over time, and use this information to move towards lower light (negative phototaxis). This work offers clear evidence for planarians using temporal sensing for navigating light gradients.

Behaviour consistent with temporal integration of light is seen while analysing multiple features of planarian gradient sensing. Animal speed and persistence of movement is sensitive to whether animals are moving away from or towards light. Animals speed up when moving away from light and vice versa. Persistence of movement is also greater. Such modulation would be consistent with animals using temporal sensing for navigation(Luo et al., 2010). Strong support for temporal sensing also comes from increased sampling as seen by larger head turns when animals are moving towards light. It would appear animals sense increasing light signal over time, and make large head turns in order to find more ‘favorable’ directions to move towards. Notably, we further show that *even in the absence of any spatial gradient (changes in light intensities over space)*, planarians are able to sense changing light intensities over time, i.e. clearly show that planarians are capable of temporal sensing (Figure 4).

The ability of planarians to use different paths while navigating their surroundings when faced with different visual conditions was previously not known. Spatial and temporal sensing have been implicated in multiple scenarios of gradient-sensing in nature, including chemotaxis and phototaxis shown by multicellular as well as unicellular organisms (Hinnemann et al., 2010; Sawin, E. P. et al., 1994). Notably, this study offers compelling evidence of highly adaptable sensory strategies and potential coexistence of spatial and temporal sensing in planarians as they navigate light stimuli. A dramatic illustration is seen with how planarians change their speed of movement while navigating light stimuli. We observe that when a spatial gradient is provided, planarians tend to slow down when moving towards light, while speeding up when moving away from light. On the other hand, the behaviour is completely inverted when no spatial gradient is provided (uniformally lit arena). When exposed to a uniformly lit arena, if light intensity increases over time (only temporal changes), worms tend to move faster. The head movement does not help the animals locate a direction of reducing intensity. This lack of feedback may cause an escape response. We suggest that animals are able to sense the complete lack of spatial gradient and move faster with increasing light intensity, as an illustration of a non-directional escape response. It is notable that in the presence of spatial gradient, the slowing down observed when moving towards light is considerably more pronounced in case of side illumination where spatial contrast is greater. Therefore, our results strongly suggests that when exposed to a spatial gradient, animals use both spatial and temporal sensing strategies. Overall, the data presented here also show that animals not only rely on both spatial and temporal sensing, but can also increase reliance on any of these sensory strategies depending on the nature of the stimuli. This is also illustrated in head-turns shown by the animals while gradient navigation. Head-turns increase as animals move towards light suggestive of temporal sensing. However, the head-turn data has distinctive features of spatial sensing when animals move perpendicular to light. When animals are oriented perpendicular to light (orientation categories C and D; Figure 1 and Figure 3), animals tend to show smaller head-turns away from light under side-illumination. In the case of top illumination, animals do tend to turn away from light but the spread of head-turn angles is larger (larger head-turns). Spatial contrast would be greater under side-illumination, reflected in the reduced sampling and more turns away from light when animals are perpendicular to light. Overall, this study highlights high degree of sensory plasticity in planarian gradient-sensing and interchanging reliance on spatial and temporal sensing.

Here, we also show a new light dependent behaviour; belly-flipping. When exposed to light from below, planarians flip their dorso-ventral orientation and attach to the top of the water surface, with the eyes facing down towards the source of light. We show this newly reported observation is mediated by the eyes. Eye-removal through RNAi abrogates this light-dependent response. Interestingly, while planarians are light aversive, this belly flipping likely indicates an active sampling and localising of the light source by the animals. Intriguingly, it has previously been reported that planarians raise their head as they sample their surroundings (Taliaferro, 1920; Walter, 1907). It was also reported that when the intensity is high enough the animals turn towards the light before orienting away potentially to identify the source(Taliaferro, 1920). It is not clear if this belly flipping behaviour is related to this previously reported behaviour of ‘head-raising’. It is conceivable that active sampling of sensory cues including light patterns may be widespread in planarians. Planarians dwell in highly variable and complex environments where light patterns and cues would be constantly evolving. Having the ability to actively sample light stimuli may be advantageous. Similarly, having the ability to simultaneously deploy multiple photosensory strategies (spatial and temporal sensing) would help animals navigate challenging and highly dynamic environments. Our results in this study support the notion of planarians having the ability to compute and navigate complex light stimuli, consistent with past evidence of complex neural processing (Shettigar et al., 2017).

Overall, this work places planarian gradient and visual light sensing well beyond a simple negatively phototactic behaviour, with refined and multi-layered sensory strategies. As mentioned above, planarians have simple prototypic eyes and possess bilateral symmetry. While much scientific effort has been focussed on better understanding complex eyes that make image forming vision possible, such simple cup-shaped non-image forming eye types can be found in a majority of the metazoan phyla(Land & Fernald, 1992). Further, simple eyes provide opportunities to link specific functional capabilities with relatively simple design features. This study significantly enriches our knowledge of functional capabilities and the rich sensory plasticity possible with a relatively simple visual system linked to a rudimentary cerebral structure. What is also highlighted is how an animal with a simple visual system uses multi-pronged strategies for gradient sensing and navigation.

While this study has not provided absolute numbers for any of the variables, analyses included here are designed to be used in a comparative manner. Planarians vary a lot in size which affects every variable(Walter, 1907). This type of analysis is meant to be used for comparing groups of animals of comparable sizes. This study offers a methodology where a systematic analysis of planarian motility and behaviour is carried out. Further, methods to analyse multiple features of planarian behaviour are developed. The variables of planarian locomotion described here can be used as a readout for other sensing-related behaviours, where even subtle phenotypes i.e. changes in planarian behaviour may be picked up. The study provides an analysis platform for various regeneration and screening studies, including RNAi and drug-screening wherein behavioural changes or changes in sensory behaviour can be analysed. This may be especially valuable for planarians that clearly show highly complex sensory and multifaceted capabilities, apart from dramatic whole body and eye-brain regeneration.

## Materials and methods

### Maintenance of planaria

Asexual strain of *Schmidtea mediterranea* were maintained in planarian media(Cebrià & Newmark, 2005) in an incubator maintained at 20 degree Celsius and fed beef liver once a week(Oviedo et al., 2008). Worms were maintained in the dark and exposed to light only during feeding and cleaning(Oviedo et al., 2008).

### Phototaxis assay

New light assays were designed (Figure 1B) using a single line of white LED lights which were wider than the circular glass arena of 120mm diameter. The LED panel was connected to a power source that could be regulated to adjust the brightness of the LEDs. The panel could be moved to shine light from the side as well as from above the arena. The light at the centre of the arena parallel to the source (Figure 1) was maintained at 50uW/cm^2^. The arena was placed on top of an IR source to allow visualisation.

The assay was conducted with 1 animal at a time for a duration of 2 minutes in a dark room maintained at 20 degrees Celsius. Before beginning the assay the animals were acclimated in this room in the dark for at least 2 hours.

The experiments with increasing and decreasing uniform illumination(Figure 5A) utilised a monitor for illumination, and a MATLAB (Mathworks, Natick, MA) program to change the intensity over time. The planarians were placed in the centre of the arena before the start of the assay, and light was changed gradually every 50ms for a duration of 90 seconds. For the experiment with eyeless animals(Figure 5C), the arena was illuminated from below and the number of animals belly up were checked at the end of the assay.

### Recording

Movies were recorded with an FLIR Grasshopper®3 High Performance USB 3.0 NIR Camera (GS3-U3-41C6NIR-C) with an IR filter placed above the arena and IR LED illumination was provided from beneath the arena for visualising the animals on the glass arena. For the uniform illumination experiments, when the illumination is from below no IR source was used and the camera was used without an IR filter. Frames were captured every 150 ms or at the framerate of 6.7 frames/sec for a duration of two minutes for the side and top illumination assays. For the uniform illumination assays the duration of recording was 90 seconds.

### RNA interference

Ovo knockdown was carried out as previously reported (Shettigar et al., 2017). The planarians were decapitated using a scalpel and ovo dsRNA was injected on the following 3 days. Control animals were injected with GFP (green fluorescent protein) dsRNA (Inoue et al., 2004). Following this treatment, the animals were allowed to regenerate for 2 weeks with media changes done every other day. The animals were screened to select those without any eyespots. These animals were then used for the phototaxis assays.

### Analysis pipeline

Custom code was written in MATLAB in order to analyse videos. The pipeline began by using a reference image of the arena taken before the experiment began to obtain the arena boundaries. This was used as a mask in the video so that only objects detected within the arena were tracked. In order to segment the worm from the background, we used a dynamic thresholding method. First, all the pixel intensities within the arena were extracted. A normal distribution was fit to the histogram of these intensity values, and the mean (mu) and standard deviation (sigma) of this fit were used to get a threshold for the frame. Most pixel values belong to the background, so we threshold the image at a value of mu - *n* * sigma. Pixels darker than this threshold are likely to be the worm. For our videos, *n* is typically 7, but it was adjusted depending on the video quality. The threshold is usually set conservatively, so the actual object is larger than the region in the binary image. Thus, after binarisation, a morphological “open” operation was performed to expand the binary region to get the true edges of the worm and any holes in the region filled. Given that the worm was always dropped in the centre of the arena, it becomes easy to identify the worm from any other spurious dark object in the frame, by considering the largest region closest to the centre of the arena in the first frame to be the worm, and then following regions closest to it in subsequent frames. Having identified the worm, we obtained the centroid using the inbuilt ‘regionprops’ function in MATLAB. No default function exists to identify the head and tail. In order to get the head and tail coordinates, the worm perimeter was first extracted and its boundary traced. Regions of maximum convex curvature are the head and tail. Since it is hard to algorithmically distinguish the two, a semi-automated approach was used. A user first confirmed whether head and tail were assigned correctly in the first frame. If not, the assignment was flipped. The head and tail in subsequent frames are those points of maximal curvature that are closest to the identified head and tail in the previous *m* frames, with *m* usually being 3. This gave us the least errors in tracking, although a user manually verified *post-hoc* that there mistakes weren’t made, and rectified them if they were. Two vectors were derived from the coordinates of the head, centroid and tail. The tail vector is the vector from the tail to the centroid, and the head vector is the vector from the centroid to the head (Supplementary Figure 1). The orientation angle and head angle were calculated from these vectors using the atan function in MATLAB. A sway was described as a head movement where the head angle is >10 degrees. The posture where the heading is within +/- 10 degrees is called a forward posture. Persistence or a run was described as the distance travelled in forward posture before a sway posture. Data for individual worms were saved and pooled for analysis *post-hoc*.

### Statistics

For the histogram fits in (Figure 2C) and (Figure 3D), a probability density function was obtained by fitting a kernel to the histogram counts of the data, using the fitdist function in MATLAB. The mode was identified as a peak in this kernel using the findpeaks function in MATLAB.

To test whether the distributions of persistent runs in the various heading directions was similar or dissimilar between the two assays, a contingency table was drawn up for the number of runs in each category, sorted by assay, following which the Pearson’s chi-squared statistic and its corresponding p-value were calculated. This was done using the crosstab function in MATLAB.

Linear mixed-effects models, results for which are reported in (Table 1) were fit in R. A maximum likelihood model was built sequentially, adding one variable to the model at a time. For example, in the case of (Figure 3C), when comparing the head angles between the “side” and the “top” assay, the assay was considered to be the fixed effect, while the identity of the worm was taken to be a random effect. The base model was one where the head angles were fit to the mean (Head Angle ∼ 1). Next, random effects were accounted for, i.e. Head Angle ∼ 1|Worm identity. Lastly, the fixed effect was added to the model: Head Angle ∼ Assay|Worm identity. To test whether “Assay” was a good predictor of the “Head Angle”, the fit models were compared using an ANOVA at a significance level of 0.01. Other linear mixed effects models were also fit and assessed using a similar, sequential model-fitting strategy.

**Table 1:** Linear mixed-effects model statistics to compare whether worms heading towards (Cat A+Cat E) and worms heading away from the light (Cat B+Cat E) had different head angles in each of the two assays - side illumination(SI) (above) and top illumination(TI) (below). Model 1 is the base model (Head Angle ∼ 1), Model 2 accounts for random effects (Head Angle ∼ 1|Worm identity). Model 3 then adds the categories as predictors to test fixed effects (Head Angle ∼ Direction|Worm identity). df - degrees of freedom; AIC - Akaike Information Criterion; BIC - Bayesian Information Criterion; L. Ratio - likelihood ratio.

|  | Model | df | AIC | BIC | logLik | Test | L.Ratio | p-value |
| --- | --- | --- | --- | --- | --- | --- | --- | --- |
| A+F vs B+E (SI) | 1 | 2 | 274460.6 | 274477.4 | -137228 |  |  |  |
|  | 2 | 3 | 272163.7 | 272188.9 | -136079 | 1 vs 2 | 2298.93 | <.0001 |
|  | 3 | 4 | 272045.3 | 272079.0 | -136019 | 2 vs 3 | 120.35 | <.0001 |
| A+F vs B+E (TI) | 1 | 2 | 283361.5 | 283378.3 | -141678.7 |  |  |  |
|  | 2 | 3 | 280597.8 | 280623 | -140295.9 | 1 vs 2 | 2765.70 | <.0001 |
|  | 3 | 4 | 280433.9 | 280476 | -140211.9 | 2 vs 3 | 167.90 | <.0001 |

## Acknowledgement

The authors would like to thank AG and DP laboratory members for critical review and discussions. RRD is grateful to the Council of Scientific and Industrial Research for fellowship support and TT Narasimhan Travel Award for travel support. We would like to thank the mechanical and electronic workshop at National Centre for Biological Sciences(NCBS)-Institute for Stem Cell Science and Regenerative Medicine(inStem) campus for their help with the assay set up. We thank the University of Hyderabad(UoH), School of Life Sciences and Department of Biochemistry for continuing support. Research in A.G. laboratory in inStem was also supported by Scientific and Engineering Research Board, Department of Science and Technology, Government of India (EMR/2017/005093). We thank Department of Biochemistry, School of Life Sciences (SLS) and UoH for support, and DBT, India for DBT-SAHAJ/BUILDER grant # BT/INF/22/SP41176/2020 to SLS. We would like to thank inStem, and the Department of Biotechnology(DBT), India for support. We would like to acknowledge Sayan Biswas for his help writing the code for the uniform illumination experiment. We would like to thank Nishan Shettigar and Anirudh Chakravarthy for discussions and valuable advice.

## Author contributions

This study was carried out under the supervision of A.G. who supplied the required materials and reagents. R.R.D. and A.G. conceived and designed the project. R.D. performed the experiments, analyzed the data, and wrote the initial manuscript. A.V. wrote the codes required for analysis of the videos and statistics and supported with reviewing the manuscript. A.N.A. assisted R.R.D. in performing behavioral experiments and experimental setup. D.P. provided expertise in planarian biology, resources, and input towards experimental design. A.G. directed the study and provided critical input, and wrote the manuscript with contributions from the other authors.

## Data and materials availability

All relevant data needed to evaluate the conclusions in the paper are present in the paper and/or the Supplementary Materials. Additional data related to this paper may be requested from the authors.

## Supplementary Material

**Supplementary Figure 1.**
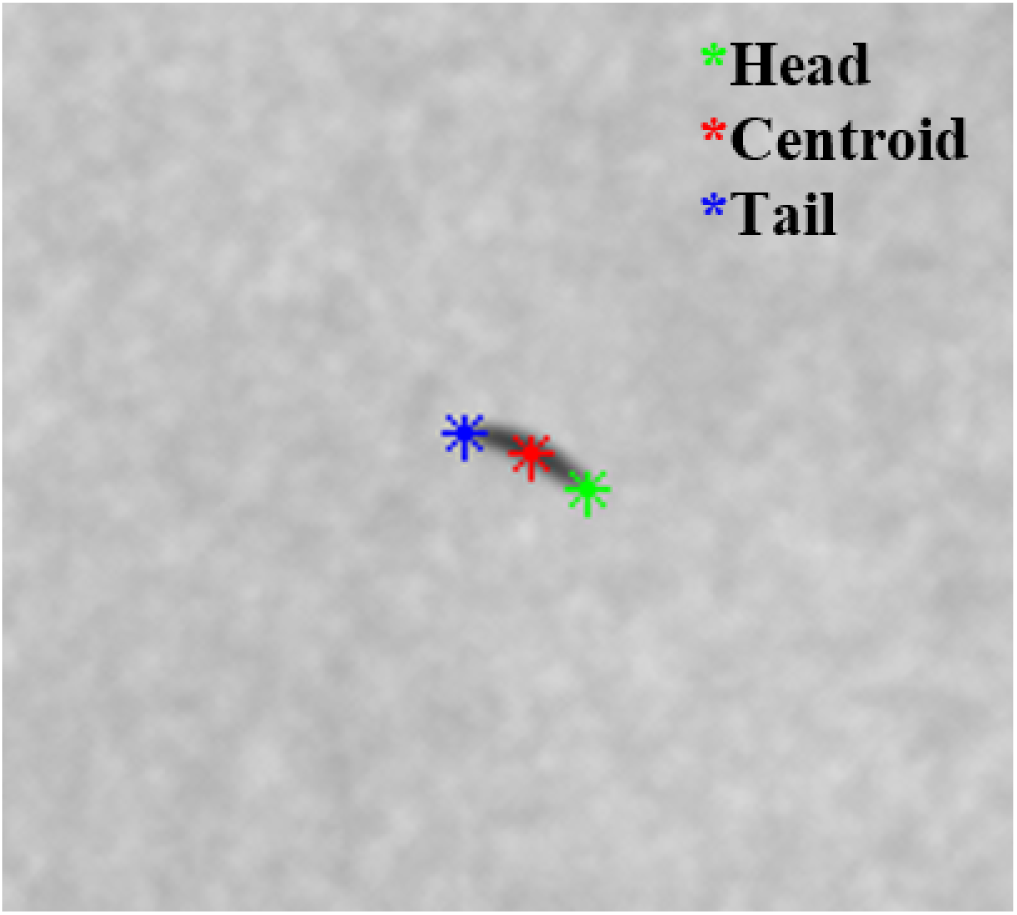
The object(worm) is identified and the head, centroid and tail is assigned as shown in this image and these points are tracked.

